# Distinctive properties of cones in the retinal clock and the circadian system

**DOI:** 10.1101/2020.09.15.297879

**Authors:** Cristina Sandu, Prapimpun Wongchitrat, Nadia Mazzaro, Catherine Jaeger, Hugo Calligaro, Jorge Mendoza, David Hicks, Marie-Paule Felder-Schmittbuhl

**Author notes:** **Corresponding authors :** MP Felder-Schmittbuhl, C Sandu. Equal contribution (Cristina Sandu, Prapimpun Wongchitrat). **Author Contributions:** CS, PW, NM, CJ, HC, JM performed experiments, analyzed data; CS, MPFS wrote the draft; DH, MPFS obtained funding; all authors reviewed the manuscript. **Competing Interest Statement:** The authors declare no potential competing interests with respect to research, authorship, and publication of this article.

## Abstract

The circadian system is a hierarchical network of cell and tissue-specific oscillators synchronized to the environmental light/dark cycle. Entrainment is mediated by the retina through diverse photosensitive cells and pigments. The retina is itself a complex circadian system whose cellular constitutive elements have not been fully characterized. By using the *Nrl*^*-/-*^ cone gain-of-function mouse model we here show that the retinal network comprises an autonomous oscillator harbored in cones. We further provide novel evidence for an input from cones to the master clock, as revealed under constant light condition.

## Introduction

Adaptation of behavior and physiology to the 24 h light/dark (LD) cycle is one of the main constraints affecting living organisms. Such adaptation is mediated by the circadian system, a network of tissue/cell-specific oscillators with an internal period close to (*circa*) 24 h, which in mammals is coordinated by the hypothalamic suprachiasmatic nuclei (SCN) (1). The retina plays a particular role in the mammalian circadian system because it is the unique photosensory input ensuring entrainment of the SCN to the LD cycle. The retina also constitutes a complete circadian system, with autonomous rhythms, resetting by light and biological outputs (2). Given the complexity of the retinal tissue comprising glial cells and six major types of neurons, how the latter contribute to retinal clock function is so far unknown. Analysis of clock gene expression *in vitro* and *ex vivo* suggested that the retina is composed of several layer-specific, coupled oscillators (3, 4). Contribution of cones to the clock network was suggested by data in chicken (5) and fish (6, 7) but has been difficult to investigate in mammals (8, 9), notably because of the scarcity of this cell type (<3% photoreceptors in mice) (10). Furthermore, importance of cones in circadian entrainment has also been somehow overlooked, in part because intrinsically photosensitive retinal ganglion cells (ipRGCs) are known as the (unique) anatomical retina-SCN link (11). Here we take advantage of the *Nrl* knock out (KO), a cone-gain of function model with all rods replaced by S-cones (12), to investigate the role of cones in the murine circadian system without the interference from rods.

## Results and discussion

### A functional clock in cones

We first characterized the cone molecular clock over 24 h in microdissected photoreceptors isolated from *Nrl* KO mice maintained in constant darkness (DD) (Figure 1A,B). Expression levels were significantly rhythmic for *Bmal1, Per1, Per2, Per3, Rev-Erb*_α_, with opposite phases between transcripts of *Bmal1* and *Per* genes. We also found significant rhythmic expression for several putative targets of the circadian clock expressed in S-cones (12): *S-opsin, Arrestin 3* and *Cnga3* (Figure 1B).

**Figure 1:**
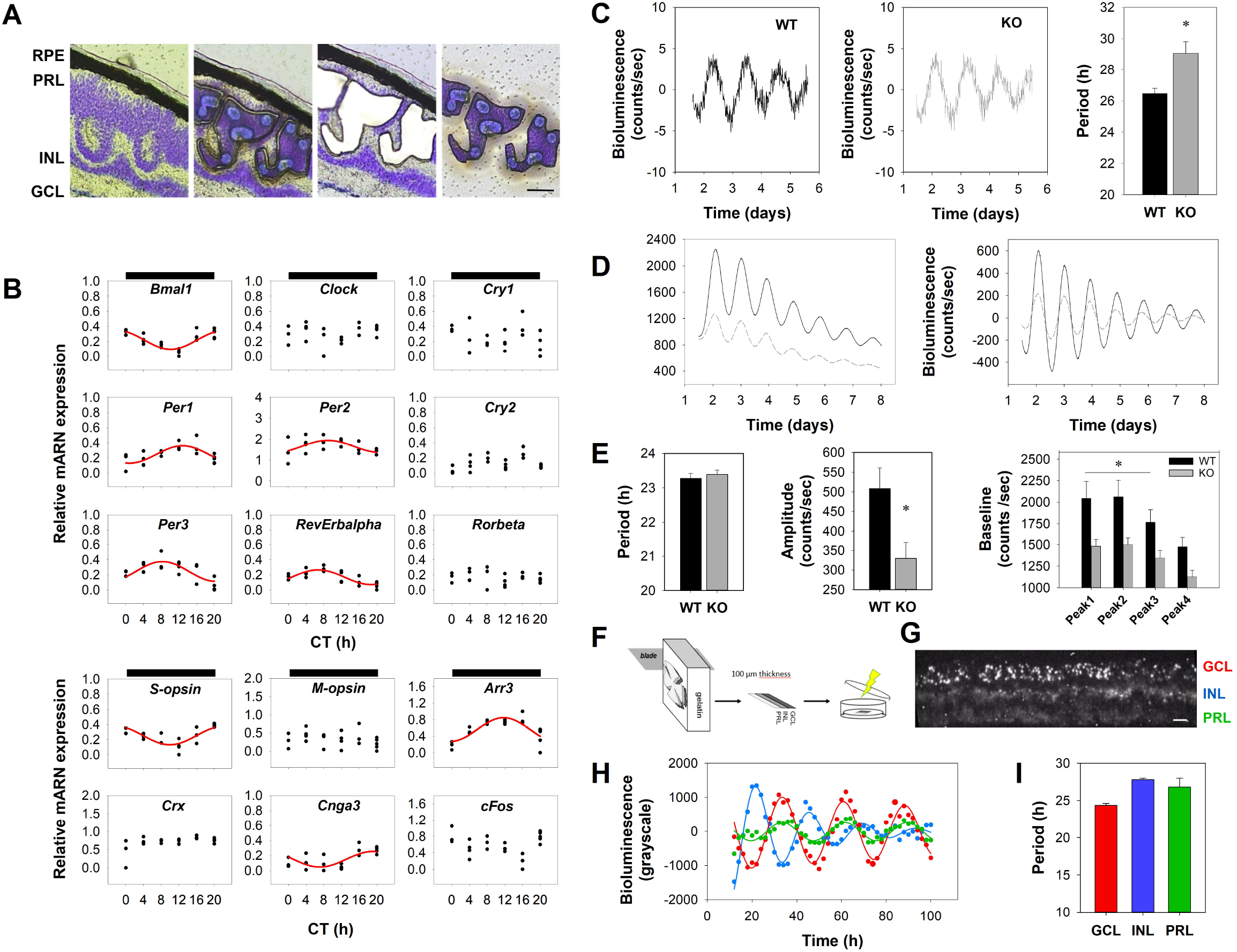
A circadian clock in cones. (A) Cresyl-violet stained retinal section showing the cone photoreceptor laser-capture microdissection from *Nrl*^*-/-*^ mice (GCL, ganglion cell layer; INL, inner nuclear layer; PRL, photoreceptor layer; RPE, retinal pigmented epithelium; scale bar, 50 µm). (B) Circadian expression profiles of clock genes (*top*) and clock output genes (*bottom*) from microdissected cone layers. Red traces show best-fitted sinewave function for significant cosinor analysis (C) Representative PER2::LUC bioluminescence recordings of cone layers isolated from WT and KO mice. Period of oscillations was significantly longer in mutants (WT n = 6, KO n = 9; *p* = 0.018). (D) Representative raw (*left*) and baseline-subtracted (*right*) bioluminescence recordings from WT and KO whole retinas. (E) When compared to WT, mutant retinas showed similar period but reduced amplitude (*p* = 0.013) and baseline levels (*p* = 0.015) (n = 12). (F) Schematic presentation of the vibratome-based strategy for isolating a transversal 100 µm thick section (G) Representative picture showing bioluminescence emission in the 3 cellular layers from a *Nrl*^*-/-*^ section (scale bar, 100 µm). (H) Representative bioluminescence counts from the 3 layers in one *Nrl*^*-/-*^ sample. Damped sinusoids represent the best-fit to subtracted data. (I) Periods were calculated separately for each cell layer and show a significant layer effect (1-way ANOVA, n = 3, *p* = 0.037). Data are presented in the histograms as mean ± SEM.

To further evaluate the capacity of cones to sustain rhythmicity, we used a vibratome-based sectioning of the retina to isolate photoreceptor (cone-only) layers from KO mice on the *Per2*^*Luc*^ reporter background, for real-time bioluminescence recordings (Figure 1C). As we previously described (4) wild-type (WT) photoreceptor layers showed robust PER2::LUC oscillations with a 26.46 ± 0.02 h period. Cone layers from KO retinas also proved robustly rhythmic but with a significantly longer period: 29.07 ± 0.03 h, probably reflecting differences in clock machinery and associated signaling occurring in rods (97% of photoreceptors in WT) versus cones or differences in coupling strength within the respective photoreceptor populations (4). Indeed, communication through gap junctions might be reduced in the S-cone enriched photoreceptor layers of the KO, since expression of connexion 36 was not detected in this cone population in mammals (13). Period difference between photoreceptor populations was, however, not reflected in whole retina explants (Figure 1D,E). By contrast, baseline levels and amplitude of PER2::LUC were reduced in the KO, likely reflecting the difference in cell numbers (≈ 40% reduction in photoreceptors in the KO (14)).

Finally, we examined how cone layers oscillate within the context of the whole retina by performing imaging-coupled bioluminescence recording on transversal sections (Figure 1F). PER2 bioluminescence signal emerged from all layers, with higher intensity in ganglion and inner cell layers and weaker signal in the outer cone layer (Figure 1G). Moreover, the signal was rhythmic in all layers (Figure 1H, SI Appendix), with distinct periods (24.28 ± 0.26 h ganglion cell layer; 27.79 ± 0.20 h inner nuclear layer; 26.80 ± 1.19 h cone layer; Figure 1I), which agrees with our previously described model of multi-oscillatory retinal clock (4). Taken together, our results demonstrate the presence of an autonomous clock in murine cones and indicate diversity in the cell-specific signaling pathways that contribute to the retinal oscillatory network.

### Major effects of a cone gain-of-function retina on the central clock

We did not detect any difference in ipRGC numbers between KO and WT retinas (Figure 2A-C), indicating that central clock characteristics in the mutant would mainly reflect the activity of cones. Wheel-running activity (WRA) profiles were similar between WT and mutants under LD and DD (Figure 2D,E). Under DD conditions, WRA endogenous period also proved similar between WT (23.96 ± 0.10 h) and KO (23.85 ± 0.05 h) (*p* = 0.278). Accordingly, in SCN explants period, rhythmic power and phase of PER2::LUC oscillations were similar between WT and KO mice (Figure 2F,G).

**Figure 2:**
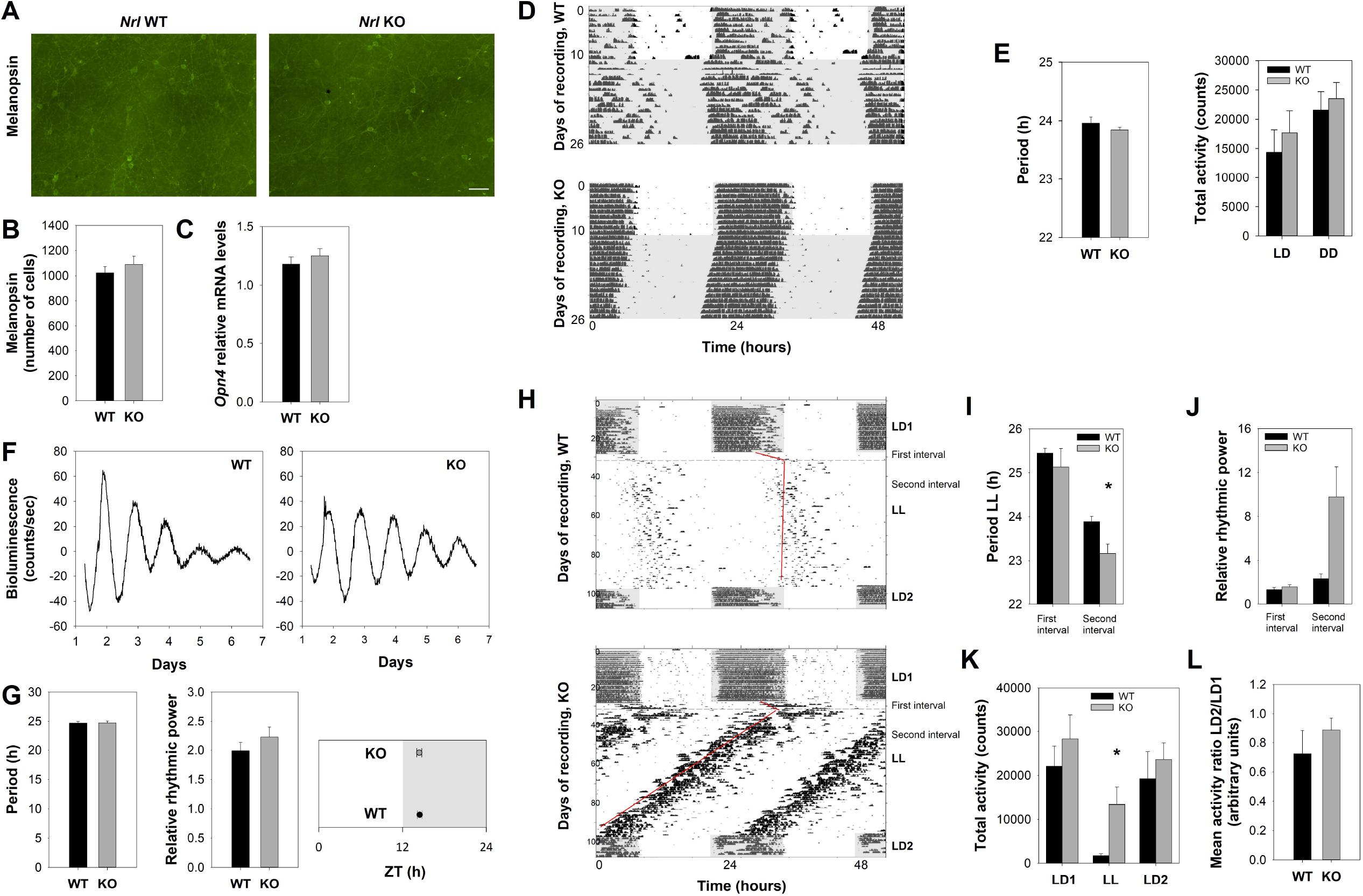
Constant light reveals the effect of cones on the central clock. (A) Immunolabelling of melanopsin in whole-mount retinas of WT and KO mice (scale bar, 50 µm). (B) Quantification of melanopsin-positive cells showing no significant difference between genotypes (WT n = 5, KO n = 6; *p* = 0.433). (C) Expression of *Opn4* transcripts in whole retina showing similar levels between WT (n = 7) and KO (n = 8) (*p* = 0.426). (D) Representative actograms of WRA of WT and KO mice in LD followed by DD (grey shading). (E) No difference between genotypes in WRA rhythms (LD, *p* = 0.579, DD *p* = 0.661), neither in endogenous period in DD (WT n = 4, KO n = 7; *p* = 0.278). (F) Representative detrended PER2::LUC bioluminescence recordings of SCN explants from WT and *Nrl*^*-/-*^ animals. (G) No genotype effect was observed in the period (*p* = 0.944), rhythmic power (p = 0.344) or phase (WT n = 5, KO n = 7; p = 0.938). (H) Representative actograms from WT and KO animals exposed to 70 days of LL (130 lux). Animals were previously maintained in LD (LD1) and returned to the same regime (LD2) following LL. Fits to onsets of activity used to determine periods are shown in red. (I) Activity pattern was separated into 2 intervals of behavior analyzed separately: a first interval with lengthened periods for both genotypes and a second one in which KO mice exhibited extremely low period values (n = 6 out of 7; *p* = 0.015). (J) Relative rhythmic power was increased in the KO during the second interval (*p* = 0.028). (K) A significant difference in total activity was found between genotypes during the second interval of LL (*p* = 0.037). Total activity did not vary for KO animals between lighting regimens (*p* = 0.066) but did for WT between LL and the LD cycles (*p* = 0.022). (L) WRA between genotypes did not differ when returned to LD (WT n = 6, KO n = 7; *p* = 0.375). Data are presented in the histograms as mean ± SEM.

To further challenge the circadian system, we exposed mice to constant light (LL). Both genotypes showed rhythmic WRA, with 2 successive steps: 1, a transient step in which periods substantially lengthen in both genotypes to approximately 25.5 h; 2, a stabilized free-run in which periods shorten, especially in the KO mice (Figure 2H,I). Thus, periods were extensively reduced in 6/7 mutants (23.17 ± 0.21 h, *p* < 0.015). Interestingly, KO mice retained robust activity, 8-fold higher than in WT (*p* = 0.035) that exhibited a drastic reduction in total WRA (Figure 2H,K). This difference was also underlined by a significantly higher rhythmic power in mutants (more than 4-fold, *p* = 0.028; Figure 2J), indicating a lack of inhibition by light and a higher robustness of circadian rhythm in KO mice. When animals were replaced in LD both genotypes showed similar phase-angle of entrainment and mean activity ratios LD2/LD1 (Figure 2H,L), confirming the integrity of light responsiveness and locomotor activity. Short free-running period values have been rarely described in LL, except in *Per2* mutants of different backgrounds (15-17). *Nrl* KO circadian phenotype in LL could be explained by distinct hypotheses: 1, sustained locomotor activity counteracts the effects of LL on neuronal activity in the SCN and counterbalances the period-lengthening mechanism (18); 2, high WRA feeds back on the clock and induces period shortening (19); 3, S-cone abundance (several hundred-fold increase with respect to WT) triggers an unknown signaling towards the central clock.

In conclusion, our study shows that cones contain an autonomous clock, contribute to the retina oscillating network and provide substantial light input to the central clock.

## Materials and Methods

Mice were handled according to the European Union Directive 2010/63/EU. Animal groups were based on previous or preliminary data and tried to conform to the 3R rule. All experimental procedures are detailed in SI.

## Acknowledgments

This work was funded by Retina France, the University of Strasbourg Institute of Advanced Studies and fellowships from Fondation Servier and ISN-CAEN to PW. We thank members of the Chronobiotron UMS 3415 for animal care and Dr. Anand Swaroop for the *Nrl* KO mice.

