## Supplementary material for "Distinctive properties of cones in the retinal clock and the circadian system": Supplentary information

[*] Equal contribution

**This PDF file includes:**

Extended methods
SI references

### Extended methods

#### Animals

*Nrl*$^{-/-}$ (C57Bl/6J background) mice were obtained from Dr. C. Grimm (Laboratory of Retinal Cell biology, University Hospital Zurich, Switzerland) with permission from Dr. A. Swaroop (NEI, Bethesda, MD, USA) (1). *mPer2*$^{Luc}$ mice (2) (C57Bl/6J background, previously purchased from The Jackson Laboratory, Bar Harbor, ME, USA) were crossed with *Nrl* mutants to generate the *Nrl*$^{-/-}$ *Per2*$^{Luc}$ and *Nrl*$^{+/+}$ *Per2*$^{Luc}$ animals. According to experiments, WT (*Nrl*$^{+/+}$) and KO (*Nrl*$^{-/-}$) animals were either homozygous for the *Per2*$^{Luc}$ knock-in allele (stated *Per2*$^{Luc}$ background) or did not contain *Per2*$^{Luc}$ allele. All mice were raised in the Chronobiotron animal facility (UMS 3415, Strasbourg, France) and housed in standard cages in groups of 3 to 4 animals, under 12h:12h light-dark (LD) cycles [ZT0-light on, ZT12-light off; broad-spectrum (400-650 nm) white light at 300 lux (MASTER PL-L 4 lamp, Philips, France); no red light at night] with food and water *ad libitum* and in an ambient temperature of 22 ± 1°C. Experiments were performed on both males and females unless otherwise stated.

#### Laser microdissection

Six weeks old *Nrl*$^{-/-}$ males (n = 30) reared in LD were exposed to constant dark (dark/dark, DD; no dim red light). After 36 h in DD mice were euthanized within the following 24 h in DD in a $CO_2$ (up to 20%) airtight chamber at the following projected ZT time points: 0, 4, 8, 12, 16, 20 (n = 5, randomly allocated, per time point). Eyes were enucleated, embedded in Tissue-Tek OCT compound (Sakura Finetek USA, Torrance, CA), frozen on dry ice and stored at −80°C until use. Animal handling and eye sampling were performed by using night vision goggles (ATN NVG-7, ATN-Optics, Chorges, France).

20 µm thick eyeball sections were cut on cryostat and placed on polyethylene naphthalate Membrane Frame slides (Life Technologies, Grand Island, NY). Three to four slides (4 sections/slide) were prepared from a single eye specimen. Each slide was stored at −80°C in a 50 mL nuclease-free tube (pre-chilled on dry ice) and used for laser microdissection within a week.

Frozen slides were thawed at room temperature for 30 s. Sections were stained with cresyl violet (1% cresyl violet acetate in 70% ethanol) for 30 s, then dehydrated through a series of ethanol solutions: 2 x 75% for 30 s, 95% for 30 s, 100% for 30 s and 100% for 2 min. Slides were air-dried at room temperature for 1 min then completely dehydrated in a vacuum chamber for 1 h before microdissection. The whole procedure was performed in RNase-free conditions.

Laser microdissection was performed using the Veritas Microdissection Arcturus system and software (Arcturus Bioscience, Inc. Mountain View, CA, USA) immediately after complete dehydration of the slides. The cone photoreceptor areas of interest were selected under microscope (20x magnification) and transferred on CapSure Macro LCM Caps (Life Technologies, Grand Island, NY) by the combined use of the infrared (power 70-80 mW, pulse 1500-3500 µs) and UV (low power 2-4) lasers (See also Figure 1A). A total cone photoreceptor area of 3 $mm^2$ was collected per eye. In order to prevent RNase reactivation and RNA degradation, the microdissection was carried out within maximum 60 min for each slide. 3-4 caps/eye were collected into the same reaction tube which contained $RLT^+$ lysis buffer (Qiagen, Hilden, Germany) and stored at −80°C.

#### Quantitative reverse-transcription PCR

Total RNA was extracted from the microdissection lysates using RNeasy Plus Micro kit (Qiagen, Hilden, Germany) according to the manufacturer's instructions and eluted in a final volume of 12 µL. RNA quantity and purity were measured using the Nanodrop ND-1000 spectrophotometer (Thermo Scientific, Wilmington, DE, USA). RNA integrity was assessed with the 2100 Bioanalyzer (Agilent Technologies, Santa Clara, CA, USA) and the RNA 6000 Pico chips (Agilent Technologies, Santa Clara, CA, USA), following the manufacturer's instructions. 25 ng of RNA from samples with the RNA integrity number (RIN) > 6 (n = 3-5) were amplified using ExpressArt mRNA amplification Nano kit (Amsbio, Oxon, UK). 150 ng of amplified RNA was reverse transcribed by using the iScript™ Advanced cDNA Synthesis Kit for RT-qPCR (Bio-Rad, Hercules, CA, USA) in a final volume of 20 µL. All samples were stored at −80°C.

Transcript levels were determined by quantitative PCR as described (3), with PCR reactions run in duplicates. The purity of the microdissected samples was verified by the absence of detection by qPCR, of transcripts for tyrosine hydroxylase (*Th*) gene and metabotropic glutamate receptor 6 (*mGluR6*) gene, as markers for the inner nuclear layer. Transcript levels were normalized to the levels of *Tbp* and *Hprt* which showed constant expression in the isolated cones over the 24 h (data not shown). All TaqMan probe-based assays were purchased from Applied Biosystems (Applied Biosystems, Foster City, CA,

USA) and designed to span exon boundaries. Data were quantified using the ΔCq method, modified to take into account gene-specific amplification efficiencies and multiple reference genes, and the qBase software (free v1.3.5) (4). log transcript levels were calculated relative to the transcript levels measured in a WT photoreceptor sample which were rescaled to one (n = 3-5 per time point). We used Excel software to detect outliers that were removed for the final statistical analysis (n = 1 for *Per1*, *Per2* and *Per3* quantification).

**Real-time bioluminescence recordings**

Bioluminescence recordings from whole retinas and isolated photoreceptor layers were obtained in several successive experiments and data were analyzed all together. Only samples generating a bioluminescence signal above the background level (similar level for both genotypes: 80 % samples for whole retinas and 30 % samples for photoreceptor layers) were retained in the study. Bioluminescence data from whole retinas and photoreceptor layers were obtained over several series of recordings.

**Whole retina explant cultures**

WT and KO mice (5-6 week-old, $Per2^{Luc}$ background), were euthanized with $CO_2$ (progressive increase up to 20% in an airtight box) during the light phase and enucleated. Eyeballs were kept at room temperature in HBSS [1 x HBSS (Sigma-Aldrich, Steinheim, Germany) containing antibiotics (100 U/mL penicillin and 100 mg/ml streptomycin, Sigma-Aldrich, Steinheim, Germany), 100 mM HEPES (Sigma-Aldrich, Steinheim, Germany) and 4.2 mM sodium bicarbonate (Sigma-Aldrich, Steinheim, Germany)] for whole retina dissection. The eyeball was incised under the ora serrata and the cornea and lens were cut out. Retinas were carefully detached from the retinal pigment epithelium and flattened with small radial incisions.

Each flattened retina was placed, photoreceptors down, onto a semipermeable membrane (Millipore, Billerica, MA, USA) in a 35 mm culture dish (Nunc, ThermoFisher, France) containing pre-incubation medium [1 ml neurobasal A medium (Gibco, Invitrogen, Life Technologies, Carlsbad, CA, USA) supplemented with antibiotics (25 U/ml penicillin and 25 mg/mL streptomycin, Sigma-Aldrich), 2% B27 (Invitrogen, Life Technologies, Grand Island, NY, USA), and 2 mM L-glutamine (Gibco, Life Technologies, Carlsbad, CA, USA). Samples were kept 24 h at 37°C in a humidified 5% $CO_2$ incubator then the medium was changed with pre-warmed (37°C) 199 recording medium [1 mL medium 199 (Sigma-Aldrich, St. Louis, MO, USA) supplemented with antibiotics (25 U/mL penicillin and 25 mg/mL streptomycin, Sigma-Aldrich), 4 mM sodium bicarbonate, 20 mM D(+)-glucose (Sigma-Aldrich), 2% B27 (Invitrogen), 0.7 mM L-glutamine (Gibco), and 100 µM beetle luciferin (Promega, Fitchburg, WI, USA)]. The medium change was performed under dim red light. Dishes were sealed with high-vacuum grease (Dow Corning; Midland, MI, USA) and placed into the LumiCycle (Actimetrics, Wilmette, IL, USA) heated at 36°C. Samples were recorded during 6-8 days and the photons were integrated for 112 s every 15 min. In bioluminescence recordings, the 2 retinas from the same animal are considered as independent, biological replicates. We here analyzed n = 12 (8 mice) for WT and n = 12 (7 mice) for KO.

**Photoreceptor layer explant cultures**

Retinas were dissected as described above. Photoreceptor layers were isolated using the vibratome technique and cultured as reported previously (5). WT (n = 6 samples, 6 mice) and KO (n = 9 samples, 8 mice) photoreceptor explants were recorded for at least 5 days and the photons were integrated for 112 s every 15 min. Exceptionally, when layers of insufficient size were collected, samples from both retinas were cultured together (2 samples in WT group, 1 sample in KO group).

**Transversal retinal slice imaging**

Flattened retinas (4 week-old KO mice, n = 3, $Per2^{Luc}$ background) were mounted with warm (37°C) 5% gelatin on top of a 10% gelatin block. The whole retina-embedded block was glued on the tissue holder and then placed into the tissue bath (containing HBSS, Sigma-Aldrich) of a Vibroslice MA752 (Campden Instruments, Loughborough, England). A transversal 100 µm thick slice was cut, placed carefully on a semipermeable membrane in a 35 mm culture dish and pre-incubated with Neurobasal A medium for 24 h. Just before imaging the medium was replaced with pre-warmed recording medium under dim red light. The sealed dish was placed into the culture chamber (37°C) of a Luminoview 200 microscope (Olympus, Hamburg, Germany) equipped with an EM-CCD camera (Hamamatsu, Japan) cooled to −76°C. Bioluminescence images (20x objective, EM gain = 80, 1 × 1 binning of pixels) were taken every 2 h over a minimum of 3 days.

#### SCN bioluminescence recordings

Animals (9 month-old, WT n = 5, KO n = 7, *Per2*$^{Luc}$ background) were killed by cervical dislocation and brains were rapidly removed and placed in ice-cold HBSS. One 500 µm coronal section of the SCN region was obtained using a stainless steel adult mouse brain slicer matrix (ZIVIC Instruments, Pittsburgh, USA), then trimmed to 1 × 1 mm. Each SCN explant (containing both nuclei) was cultured onto a Millicell culture membrane (Merck Millipore Ltd, Tullagreen, Ireland) in a 35-mm culture dish with 1 mL of DMEM (Sigma-Aldrich) supplemented with 0.35% D(+)-glucose, 0.035% sodium bicarbonate, 10 mM HEPES, 2% B27, antibiotics (25U/mL penicillin and 25mg/mL streptomycin) and 0.1 mM beetle luciferin. Culture dishes were sealed with vacuum grease. The bioluminescence was recorded using the LumiCycle for 112 s in 15 min intervals and during at least 6 days.

#### Bioluminescence data analysis

Whole retina and SCN explant PER2::LUC raw data were subtracted with a 24 h running average (removal of the baseline drift) using the LumiCycle analysis software (Actimetrics, Wilmette, IL, USA). The first cycle was removed and the analysis was performed on the following 4 (retina) or 5 (SCN) cycles. The robustness of the rhythms (relative rhythmic power (6)) and the phase were also calculated using the LumiCycle analysis software. The phase of the first peak in SCN samples was expressed relative to the LD cycle to which animals were previously exposed.

The period and amplitude were determined using a cosinor derived sine wave function: $f = y0 + a * \exp(-x/d) * \sin[2 * \pi * (x + c) / b]$ where a is the amplitude (counts/s), b is the period (h), c is the phase-related term (h) and d is the damping rate (days) and assuming that damping follows an exponential pattern. Baseline for each individual peak in retinal samples was estimated as the baseline from LumiCycle analysis taken at the peak time. Photoreceptor layer data were analyzed as previously described, on 4 successive cycles (5).

Transversal retinal images were analyzed with ImageJ (open source software https://imagej.nih.gov/). A median 3D filter was applied to remove the hotspots. The ganglion cell layer (GCL), inner nuclear layer (INL) and photoreceptor layer (PRL) were defined as regions of interest (ROI) and the bioluminescence levels (grey levels) were measured and exported for the analysis of rhythmicity. The periods were determined using the cosinor derived sine wave function: $f = y0 + a * \exp(-x/d) * \sin[2 * \pi * (x + c) / b]$ as above.

#### Locomotor activity recordings

For behavioral recordings, male and female mice (WT and KO combined with the *Per2*$^{Luc}$ knock-in allele) were housed in individual standard cages equipped with a 10-cm-diameter stainless steel running wheel (7). Data were collected in 5 min bins and analyzed with the ClockLab Software (Actimetrics, Wilmette, IL, USA). Locomotor activity data were represented as double-plotted in actograms.

#### Circadian phenotype

To determine the daily and circadian rhythm of locomotor activity in *Nrl* mutant mice, 5-6 month-old mice (WT n = 4, KO n = 7) were initially maintained for 10 days under LD 12:12 and then 16 days under constant darkness (DD). Total activity levels were calculated over the last 3 days in LD and the endogenous period (Chi-square Periodogram method) was determined over a 9-day interval, 7 days after the transition to DD.

#### Exposure to constant light

Effects of constant light exposure (light/light, LL) were assessed by wheel-running activity in 6 month-old mice (WT n = 6, KO n = 7). Thus**,** after 10 days in LD 12:12 animals were transferred to LL for 70 days (130 ± 34 lux on average). Total activity per cycle, period and relative rhythmic power were measured by using ClockLab. Mice were then exposed to a second LD cycle (LD2: 10 days) to evaluate if entrainment and locomotor activity returned to baseline levels.

#### Statistical analyses

Results are expressed as means ± SEM, except for qPCR data. Statistical analyses were performed by using SigmaPlot 12 software (Systat Software, San Jose, CA, USA). Comparison of two groups was performed by using the Student's t-test. Comparison of several groups was performed by using 1-way or 2-way ANOVA for independent and repeated measures, followed by post-hoc test (Holm-Sidak test).

Data from qRT-PCR over 24 h in DD were also analyzed by nonlinear least-square fitting of a 24 h sinusoid (cosinor analysis) $f = a + [b*\cos(2*\pi*(x - c)/24)]$ (8). A statistically significant difference was assumed with $p$ values less than 0.05.
